# Spatially localized ligand binding to receptors affects magnitude and timing of signaling response

**DOI:** 10.64898/2026.06.23.734049

**Authors:** Nicholas T. Duong, Sahar A. Kamil, Jayde Casimir-Powell, Costin N. Antonescu, Aidan I. Brown

## Abstract

Cell surface receptors are activated by ligand binding and transmit signals into the cell. Epidermal growth factor (EGF) receptor (EGFR) signaling regulates cell growth, differentiation, and survival, and its dysregulation is linked to cancer. Recent experiments show that ligand binding to EGFR is enhanced for receptors in tetraspanin nanodomains on the cell surface. We use kinetic modeling of receptor confinement, ligand binding, and internalization to compare confinement and signaling behavior for EGFR with spatially localized ligand binding to a hypothetical receptor that has uniform ligand binding anywhere on the cell surface. We find that introducing a membrane domain that confines and enhances ligand binding to receptors leads to more consistent confinement across ligand levels, raises necessary ligand levels for steady-state signaling, and flattens and extends the signaling response to sudden ligand concentration increases. This confining domain that enhances ligand binding provides the cell with a distinct regulatory mechanism to tune its signaling response. We also find that the concentration of receptors in signaling states and the fraction of receptors in signaling states respond to ligand at different ligand concentrations, with substantial increase of the concentration of receptors in signaling states occurring at a much lower ligand concentration than a substantial increase of the fraction of surface receptors in signaling states. This quantitative modeling of spatially restricted receptor activation applies to other receptors with similar characteristics and builds towards physical principles of receptor signaling.

## INTRODUCTION

Cell surface receptors facilitate the transfer of information from the extracellular to the intracellular environment, allowing cells to respond to signals carried by ligand molecules [1–4]. Epidermal growth factor (EGF) receptors (EGFR) regulate key physiological signaling processes and are involved in cell growth, differentiation, and migration [1, 2, 5, 6]. Dysregulation of EGFR signaling activity is associated with many cancers, and accordingly these receptors are a target for anticancer therapy and drug development [5, 7–9].

EGF receptors are activated by at least seven distinct ligand molecules, with different ligand types having variable binding lifetimes [10–12], indications of different capacity to induce receptor interaction with clathrin [13], and distinct signaling outcomes [14, 15]. Following ligand binding, conformational changes in EGFR drive interactions with signaling molecules that trigger phosphorylation to induce downstream signaling responses [16–18]. Ligand bound EGFR are recruited into and internalized from clathrin domains via clathrin mediated endocytosis [19–21].

Standard models describe ligand binding to a cell surface receptor as uniformly occurring at the same rate for a receptor at any cell membrane location. Recent experimental work showed enhanced ligand binding to EGFR within tetraspanin nanodomains, compared to other locations on the cell membrane [22]. Tetraspanin nanodomains are 50 – 200 nm-wide regions of the cell membrane enriched in tetraspanin proteins that scaffold the organization of other proteins [23–26]. This confinement of EGFR to tetraspanin nanodomains, which facilitates ligand binding, is earlier in the signaling process than the well-studied confinement of EGFR to clathrin domains. The signaling consequences of spatially localized ligand binding to EGFR and other cell surface receptors are not understood.

The contribution of spatial factors to cell signaling, such as localization of certain processes to specific domains, is well recognized [27], including for EGFR signaling [28]. EGFR location on the cell surface or within the cell leads to different interactions and contributes to distinct downstream signaling [29]. We build on this existing understanding of spatial contributions to cell signaling, focusing on EGFR.

For an EGFR-type receptor that is recruited to a confining domain and internalized following ligand binding, we use kinetic modeling of receptor confinement and signaling to compare spatially localized ligand binding with uniform ligand binding across the cell surface. We find that localized ligand binding (compared to uniform ligand binding) alters receptor confinement behavior to be consistent across ligand concentrations, shifts steady-state signaling to higher ligand concentrations, suppresses and extends signaling increases upon increases in ligand stimulation, and overall provides a distinct means of receptor signaling regulation.

## METHODS

To investigate the effect of spatially localized ligand binding on signaling behavior, we use quantitative models of distinct receptor monomer states and the transition rates between them. Compared to the uniform ligand binding model, the spatially localized ligand binding model introduces a tetraspanin domain state from which ligand binding is increased compared to ligand binding to receptors outside this state.

We describe cell surface states for EGF receptors with spatially localized ligand binding with a four-state kinetic model (Fig. 1A), which we term the localized model. For this model, unconfined receptors without bound ligand (concentration *R*_S_) arrive on the cell surface at rate *k*_p_. Unliganded, unconfined receptors are internalized at a background rate *k*_bi_, are recruited to tetraspanin nanodomains at rate *k*_+T_, and bind ligand at rate *k*_+L,slow_. Receptors in a tetraspanin domain (*T*_S_) exit the domain without binding ligand at rate *k*_-T_ and bind ligand and exit the domain at rate *k*_+L_. Unconfined ligand-bound receptors (*R*_L,S_) unbind ligand at rate *k*_-L_ and are recruited to clathrin domains at rate *k*_+c_. Receptors in a clathrin domain (*R*_c,S_) exit the domain at rate *k*_-c_ and become internalized at rate *k*_i_. Noting the two ligand-binding pathways in the localized model, the enhanced ligand binding for receptors in tetraspanin domains is expressed as *k*_+L_ = 10*k*_+L,slow_.

**FIG. 1.**
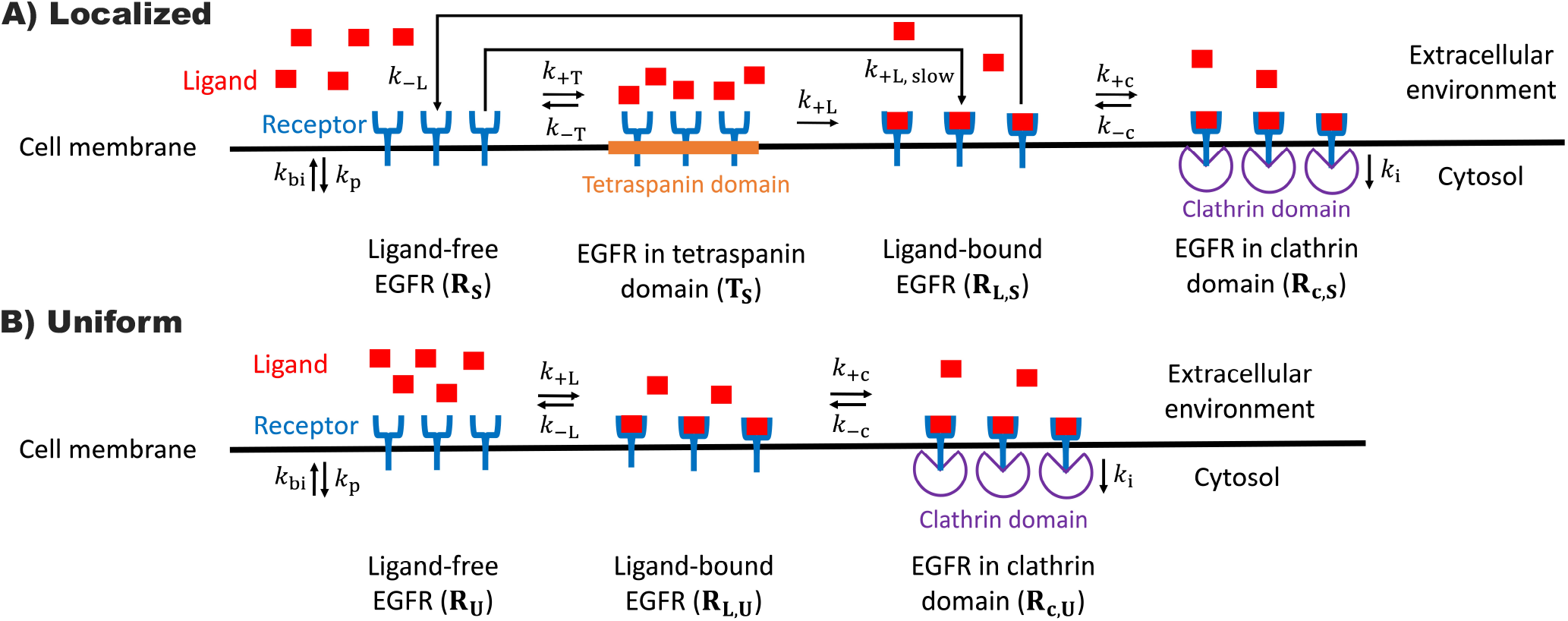
Kinetic models of EGFR dynamics on the cell surface. Receptors (blue) occupy discrete states on the cell surface with transition rate consants *k* that are summarized in Table 1. A) The spatially localized ligand binding model enhances ligand (red squares) binding in tetraspanin domains *T*_S_ (orange region). Unliganded, unconfined receptors *R*_S_ arrive on the surface at rate *k*_p_. These unliganded, unconfined receptors are recruited to tetraspanin domains (rate *k*_+T_), become ligand-bound (*k*_+L,slow_), and are internalized (*k*_bi_). Receptors in tetraspanin domains *T*_S_ become ligand bound (*k*_+L_) or exit the domain (*k*_-T_). Ligand-bound receptors *R*_L,S_ are recruited to clathrin domains (purple wedged open circle) (*k*_+c_) or the ligand unbinds (*k*_-L_). Receptors in clathrin domains *R*_c,S_ are internalized (*k*_i_) or exit the domain (*k*_-c_). Ligand binding to receptors in tetraspanin domains (*k*_+L_) is 10-fold faster than ligand binding to unliganded receptors outside of tetraspanin domains (*k*_+L,slow_). B) The uniform ligand binding model involves similar unliganded *R*_U_, ligand-bound *R*_L,U_, and clathrin-confined *R*_c,U_ receptor states as (A), without the tetraspanin state and associated transitions. The rate of ligand binding to *R*_U_ is the same as the rate of ligand binding to *T*_S_ in (A).

We describe EGF receptors with uniform ligand binding with a similar three-state kinetic model (Fig. 1B), which we term the uniform model, that omits the tetraspanin domain (*T*_S_) state and associated transitions, but with other states (*R*_U_, *R*_L,U_, *R*_c,U_) analogous to the localized model. The uniform ligand binding model also increases the ligand binding rate *k*_+L,slow_ for un-confined receptors without bound ligand (*R*_U_) to *k*_+L_, such that the sole ligand binding pathway of the uniform ligand binding model has an equal rate to the primary ligand binding rate of the spatially localized ligand binding model, to provide a direct comparison. Differential equations describing both the spatially localized ligand binding model and uniform ligand binding model, and their steady-state solution, are provided in the Appendix. The parameters for both models and their estimated values are summarized in Table I, with the rationale for parameter value estimates provided in the Appendix.

**TABLE I.** Model parameters and estimated values. The value of *k*_p_ is arbitrary and sets the scale of receptor numbers. *k*_+L_ and *k*_+L,slow_ vary with ligand concentration. The estimated ligand unbinding rate *k*_-L_, shown for EGF, varies between ligand types.

| Parameter | Description | Estimated rate constant |
| --- | --- | --- |
| $k_p$ | Cell surface arrival | $1 \text{ s}^{-1}$<br>(arbitrary) |
| $k_{bi}$ | Background internalization | $3 \times 10^{-4} \text{ s}^{-1}$ |
| $k_{+T}$ | Tetraspanin domain entry<br>*Localized model only* | $0.033 \text{ s}^{-1}$ |
| $k_{-T}$ | Tetraspanin domain exit<br>*Localized model only* | $0.12 \text{ s}^{-1}$ |
| $k_{+L}$ | Primary ligand binding | Varies |
| $k_{+L,slow}$ | Slow ligand binding | $0.1 k_{+L}$ |
| $k_{-L}$ | Ligand unbinding | $0.05 \text{ s}^{-1}$<br>for EGF |
| $k_{+c}$ | Clathrin domain entry | $0.1 \text{ s}^{-1}$ |
| $k_{-c}$ | Clathrin domain exit | $0.05 \text{ s}^{-1}$ |
| $k_i$ | Internalization | $0.05 \text{ s}^{-1}$ |

A key estimate is the relationship between ligand concentration and the primary ligand binding rate to a receptor *k*_+L_. We estimate that EGF concentration in units of ng/mL is 50× the rate of ligand binding to a receptor in units of s^−1^ using diffusion-limited rates (see Appendix), which assumes that all receptors are able to bind ligand for the primary binding pathways. With typical physiological EGF concentration from 0.2 – 0.7 ng/mL [30–32], a typical ligand binding rate is 0.01 s^−1^ (corresponding to 0.5 ng/mL); or for elevated EGF concentration of 5 – 10 ng/mL [22, 33] the binding rate is 0.1 s^−1^ (corresponding to 5 ng/mL).

## RESULTS

### Confinement behavior

We use these models to calculate the fraction of EGF receptors confined to tetraspanin and clathrin domains for the localized model and to clathrin domains for the uniform model. The fraction *f*_c_ of confined receptors for the localized and uniform models, respectively, are

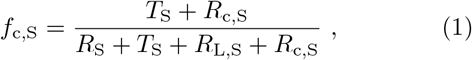

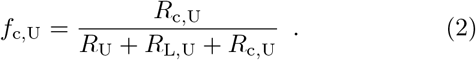

Figure 2A, which shows the confined fractions *f*_c,S_ and *f*_c,U_ across a wide range of ligand binding rates, illustrates that the localized model has a nonzero confinement fraction at a ligand binding rate of zero, in contrast with the uniform model which has confinement of zero at a ligand binding rate of zero. Experimental data shows that, across EGF concentrations from zero up to high physiological concentrations of 50 ng/mL, approximately 20% of EGF receptors are not mobile [22]. Thus the uniform ligand binding model is inconsistent with the experimental data because the confinement level for the uniform model is nearly zero for low ligand binding rates.

**FIG. 2.**
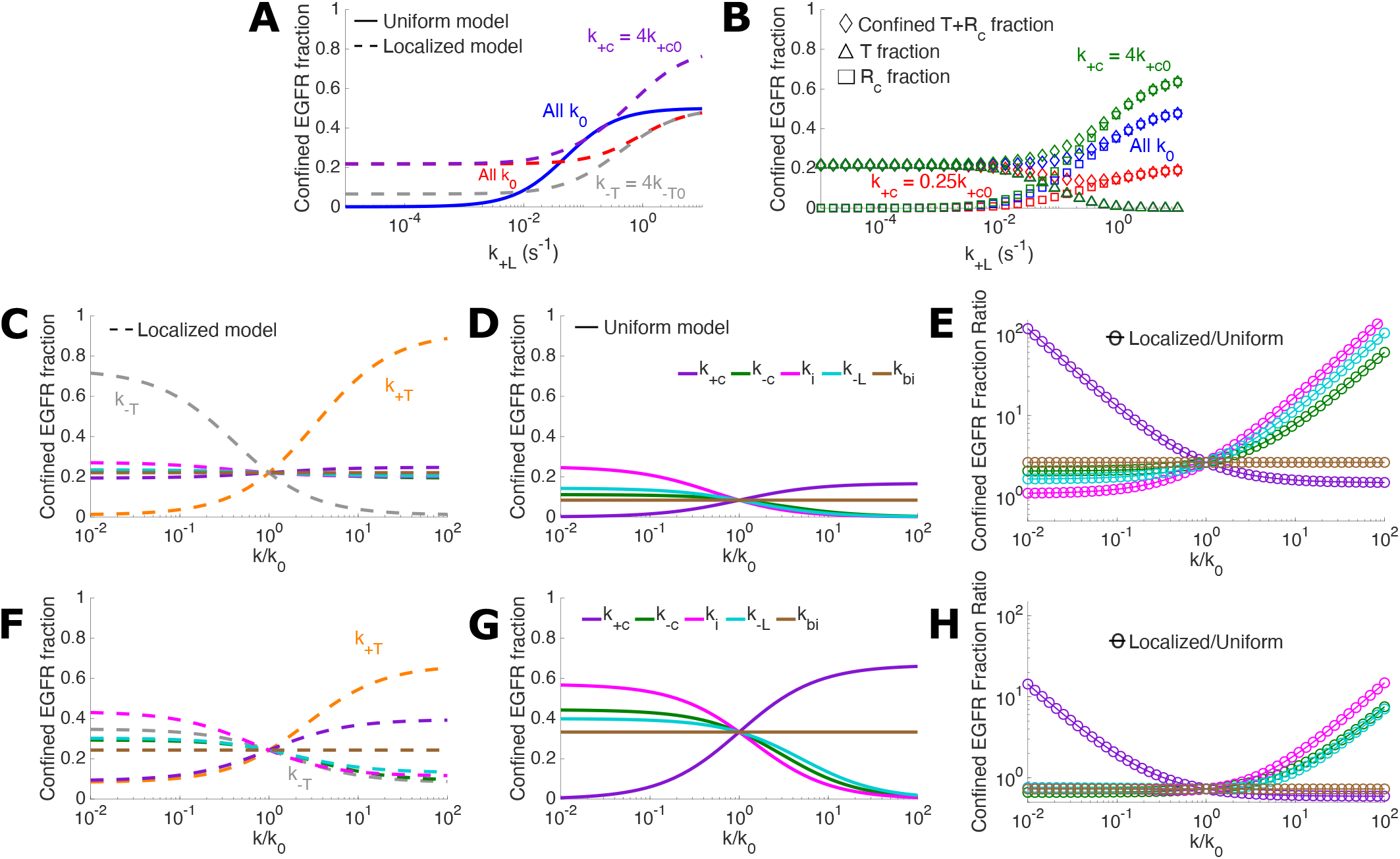
EGFR confinement behavior. (A) Fraction *f*_c_ of EGF receptors confined to tetraspanin and clathrin domains as ligand binding rate *k*_+L_ is varied. *k*_+c0_ and *k*_-T0_ are the estimated parameter values from Table I. *k*_+c_ and *k*_-T_ varied as indicated by curve color. (B) For the localized model, fraction of EGF receptors confined to both tetraspanin and clathrin domains (diamonds), tetraspanin domains (triangles), and clathrin domains (squares) as ligand binding rate *k*_+L_ is varied. Point color indicates parameter values. (C,D) Confined EGFR fraction *f*_c_ as parameters are individually varied for ligand binding rate *k*_+L_ = 0.01 s^−1^. (C) is for the localized model and (D) is for the uniform model. (E) Ratio *f*_c,S_*/f*_c,U_ of confined receptor fraction for the localized model to the uniform model as the values of the parameters shared between the two models are varied, for ligand binding rate *k*_+L_ = 0.01 s^−1^. (F,G) Confined EGFR fraction *f*_c_ as parameters are individually varied for ligand binding rate *k*_+L_ = 0.1 s^−1^. (F) is for the localized model and (G) is for the uniform model. (H) Ratio *f*_c,S_*/f*_c,U_ of confined receptor fraction for the localized model to the uniform model as the values of the parameters shared between the two models are varied, for ligand binding rate *k*_+L_ = 0.1 s^−1^. For panels C – H, *k/k*_0_ indicates the ratio of the parameter value and the estimated parameter value listed in Table I and the legend in D indicates the curve color for parameter values shared by both models. For panels C and F, the legend in C indicates the curve color for the tetraspanin entry and exit rates. For A, C, D, F, and G, solid curves are for uniform model and dashed curves are for localized model. All parameter values are those listed in Table I unless otherwise specified. Confined fractions *f*_c_ are from Eqs. 1 and 2.

The upper end of the EGF concentration range in experiments with 20% of EGFR not mobile is approximately 50 ng/mL, which corresponds to a 1 s^−1^ ligand binding rate. With our estimated parameters, the localized model has a confined receptor fraction of approximately 20% until the binding rate increases to nearly 1 s^−1^, consistent with experimental measurements. The localized model also predicts that the confined receptor fraction will double for ligand binding rates exceeding approximately 5 s^−1^ (250 ng/mL) — experimental data shows that the confined fraction of receptors begins to rise for EGF concentrations of 100 ng/mL [22].

Figure 2A shows that variation of different parameters affects the confined fraction of receptors. For the localized model, varying the tetraspanin exit rate *k*_-T_ changes the confined receptor fraction at low ligand binding rate but not at high ligand binding rate because at low ligand binding rate the receptor confinement is primarily in tetraspanin domains and at high ligand binding rate confinement is primarily in clathrin domains. Conversely varying the clathrin domain entry rate *k*_+c_ changes the confined receptor fraction at high ligand binding rate but not at low ligand binding rate. For the localized model, Fig. 2B decomposes the confined receptor fraction into tetraspanin and clathrin domain contributions. This decomposition shows that at low ligand binding rate the confined receptors are in tetraspanin domains but absent from clathrin domains, and at approximately a ligand binding rate of 10^−2^ s^−1^ the confined receptors begin a switch between these domains, such that at sufficiently high ligand binding rate the confined receptors are in clathrin domains and absent from tetraspanin domains. The physiological EGF concentration range (corresponding to approximately 0.5 – 10 ng/mL, or ligand binding rates of 0.01 – 0.5 s^−1^) covers a range with substantial confinement to both tetraspanin and clathrin domains.

The uniform model has zero receptor confinement at low ligand binding rate because there is no domain that confines receptors prior to ligand binding. Once receptors are ligand bound, they may be recruited to and confined within clathrin domains until internalization or ligand unbinding. The localized model has consistent receptor confinement across ligand binding rates because the lower ligand binding rates are insufficient to substantially remove receptors from tetraspanin domains and higher ligand binding rates have a decrease in confinement to tetraspanin domains that is approximately balanced by an increase in confinement to clathrin domains. At sufficiently high ligand binding rates (≥ 1 s^−1^, corresponding to EGF concentrations ≥50 ng/mL) the receptors are largely absent from tetraspanin domains and the confinement level is controlled by the extent to which ligand-bound receptors are confined to clathrin domains.

At a lower physiological ligand concentration (ligand binding rate of 10^−2^ s^−1^), Fig. 2C shows how the confined fraction of receptors is affected by individually changing each parameter of the localized model from its estimated value without changing other parameter values. Most parameters have limited effect on the confined receptor fraction, with the exception of the entry rate *k*_+T_ and exit rate *k*_-T_ from tetraspanin domains. Increasing (decreasing) *k*_+T_ substantially increases (decreases) the confined receptor fraction, and increasing (decreasing) *k*_-T_ substantially decreases (increases) the confined receptor fraction. This control of receptor confinement by tetraspanin domain entry and exit rates is because at this ligand binding rate most confined receptors are within tetraspanin domains, and few receptors are ligand-bound.

Similar to Fig. 2C for the localized model, Fig. 2D shows how the confined fraction in the uniform model is affected by parameter changes. For the uniform binding model, all confined receptors are in clathrin domains. All parameters except the background internalization rate *k*_bi_ can substantially affect the confined receptor fraction: a lower clathrin entry rate *k*_+c_, higher clathrin exit rate *k*_-c_, or higher internalization rate *k*_i_ substantially decreases receptor confinement; and higher *k*_+c_ or lower *k*_i_ substantially increases receptor confinement.

Figure 2E compares receptor confinement between the localized and uniform models as the parameters shared by the two models are individually varied, showing the localized model, with additional tetraspanin confining domain, has consistently higher confinement (data above one on the vertical axis). Lower clathrin entry rate *k*_+c_, higher clathrin exit rate *k*_-c_, higher ligand unbinding rate *k*_-L_, and higher internalization rate *k*_i_ lead to much higher confined receptor fractions for the spatially localized model because these parameter ranges lead to few receptors in clathrin domains in both models, such that there are comparatively many receptors in tetraspanin domains in the spatially localized model.

Figures 2F-H show how the confined EGFR fraction changes as the parameter values are changed with a higher ligand binding rate of *k*_+L_ = 0.1 s^−1^ corresponding to higher physiological EGF concentration of approximately 5 ng/mL. Figures 2F and 2G show that at this higher ligand binding rate the rates of tetraspanin domain entry and exit (*k*_*±*T_) have less effect on confinement than other parameters (*k*_*±*c_, *k*_-L_, *k*_i_), compared to the lower ligand binding rate in Figs. 2C and 2D. This lower confinement sensitivity to tetraspanin domain parameters and higher sensitivity to other parameters is because at higher ligand binding rate fewer of the receptors are confined to tetraspanin domains and more are confined to clathrin domains (Fig. 2B). For the estimated parameter values, at this higher ligand binding rate the localized model has approximately one-third less confinement than the uniform model (Fig. 2H). Increasing *k*_-c_, *k*_-L_, and *k*_i_ or decreasing *k*_+c_ allows the localized model to have more confinement than the uniform model (Fig. 2H).

### Steady state signaling

Ligand binding and clathrin localization both contribute to EGFR signaling [26, 34–36]. We represent the EGFR signaling level by the sum of the concentrations of EGFR in ligand-bound and non-clathrin localized (*R*_L_) and ligand-bound and clathrin-localized (*R*_c_) states, which we will refer to as signaling states, assuming that other signaling aspects such as receptor phosphorylation or kinase activity will be downstream of the ligand binding and clathrin localization status of the receptors. We use our model to evaluate how spatially localized ligand binding affects the population of EGFR in these signaling states. We initially focus on steady state signaling, when the receptor populations have stopped changing during periods of consistent ligand stimulation.

Figure 3A shows how concentration of receptors in signaling states states responds to increasing ligand binding rate (proportional to ligand concentration). At sufficiently low ligand binding rate very few receptors are in a signaling state. With increasing intermediate ligand binding rate the concentration of receptors in signaling states increases. At high ligand binding rate the signaling receptors plateau at a high level. The uniform model has a substantial concentration of receptors in signaling states at a lower ligand binding rate than the localized model. This shift is because the uniform model allows all receptors without ligand bound to bind ligand at a high rate, but the localized model only binds ligand to receptors in tetraspanin domains at a high rate, such that the localized model requires a higher ligand binding rate to reach the same level of receptors in signaling states.

**FIG. 3.**
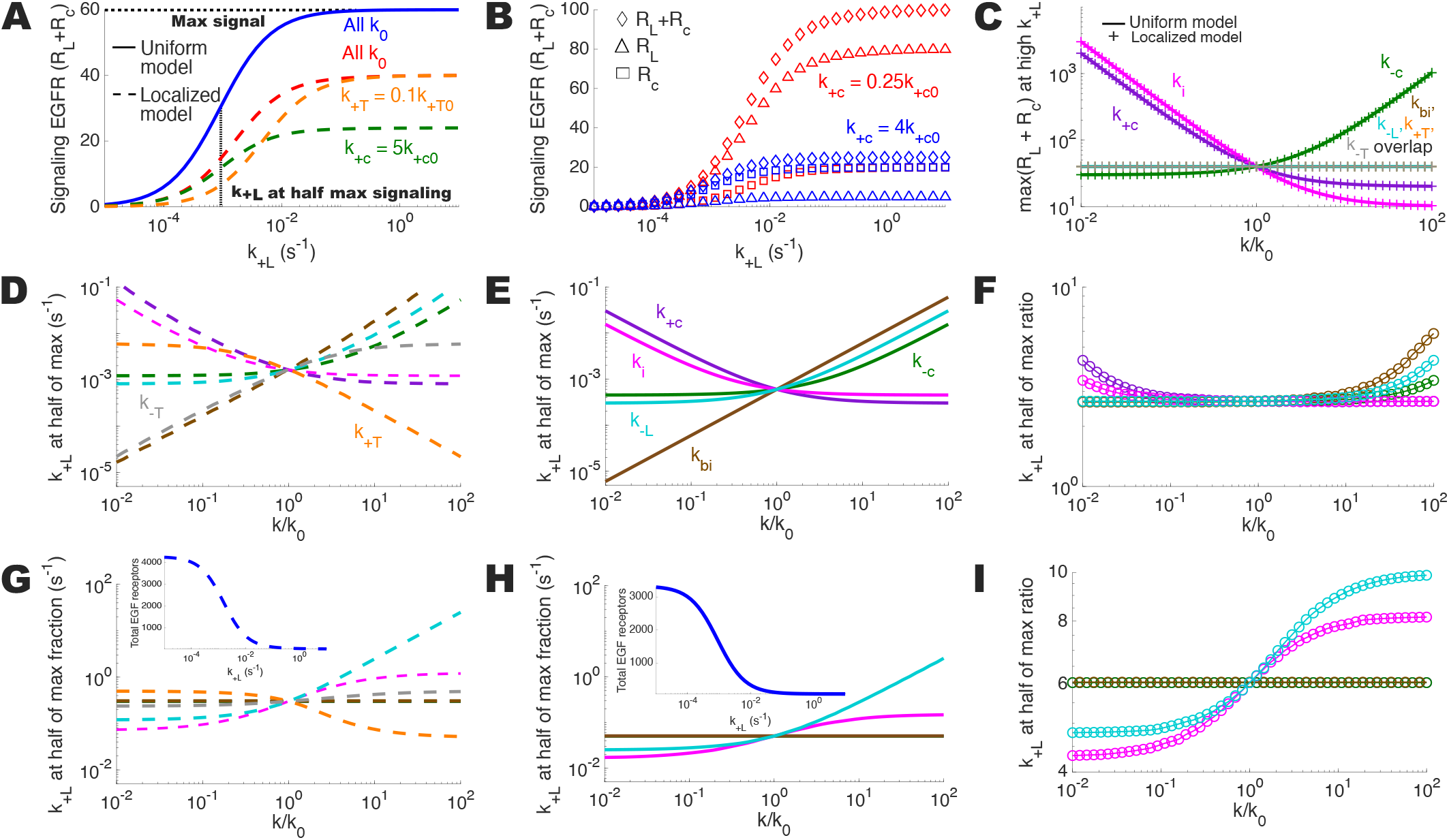
EGFR steady state signaling behavior. (A) Signaling EGF receptor (*R*_L_ + *R*_c_) concentration as the ligand binding rate is varied. Parameter values indicated by curve color, with ‘All *k*_0_’ indicating that all model parameters are the estimated parameter values from Table I, and *k*_+T0_ and *k*_+c0_ referring to the individual estimated parameter values from Table I. The horizontal dotted black line indicates the maximum level of signaling receptors and the vertical dotted black line indicates the ligand binding rate *k*_+L_ at which the level of signaling receptors is half the maximum level of signaling receptors, both for the blue solid uniform model curve. (B) Signaling EGF receptor (*R*_L_ + *R*_c_, diamonds), ligand-bound and non-clathrin localized receptor (*R*_L_, triangles), and ligand-bound clathrin-localized receptor (*R*_c_, squares) concentration as the ligand binding rate is varied. Localized model results. (C) Maximum signaling EGF receptor concentration (illustrated by horizontal dotted line in panel (A)) as parameters are individually varied. *k/k*_0_ indicates the ratio of the parameter value and the estimated parameter value listed in Table I. (D,E) Ligand binding rate at which *R*_L_ + *R*_c_ reaches half of its maximum (illustrated by vertical dotted line in panel (A)) as parameters are individually varied for (D) the localized model and (E) the uniform model. (F) Ratio, of the localized model to the uniform model, of the ligand binding rate *k*_+L_ at which the signaling receptor concentration *R*_L_ + *R*_c_ reaches half of its maximum. (G,H) Ligand binding rate at which the fraction of receptors in the signaling states *R*_L_ and *R*_c_ reaches half of its maximum as parameters are individually varied for (G) the localized model (Eq. 3) and (H) the uniform model (Eq. 4). Insets show the total receptor concentration (sum of all states) as the ligand binding rate is varied. (I) Ratio, of the localized model to the uniform model, of the ligand binding rate *k*_+L_ at which the signaling receptor fraction reaches half of its maximum. Curve color labels in panel (E) also apply to other panels (D)–(I) and curve color labels in (D) also apply to panel (G). For panels A, D, E, G, and H, solid curves are for the uniform model and dashed curves are for the localized model. All parameter values are those listed in Table I unless otherwise specified.

For the localized model, Fig. 3A illustrates the distinct effect of different parameters. Increasing the clathrin recruitment rate *k*_+c_ decreases the level of signaling receptors at saturating ligand binding rate without changing the ligand binding rate at which the signaling receptor level rises — this decrease in saturated signal receptor level is because receptors in clathrin domains are removed from the cell surface. In contrast, decreasing the tetraspanin recruitment rate *k*_+T_ increases the ligand binding rate *k*_+L_ required for the signaling state concentration to rise without changing the level of signaling receptors at saturation — this increase in ligand binding rate is because there are fewer receptors in tetraspanin domains that can have ligand bind at a high rate.

For the localized model, Fig. 3B shows the decomposition of the receptor signaling states into ligand bound and non-clathrin localized (*R*_L_) and ligand-bound and clathrin-localized (*R*_c_) states for varying clathrin domain entry rates. At relatively low clathrin entry rate *k*_+c_, more receptors are in signaling states, with most of these receptors in the ligand-bound non-clathrin localized state (*R*_L,S_) and relatively few in the ligand bound clathrin-localized (*R*_c,S_) state. This pattern at low *k*_+c_ is due to receptors entering clathrin slowly and thus not being as exposed to removal from the cell surface via internalization, leading to a higher receptor population. At relatively high *k*_+c_, fewer receptors are in signaling states, with most of these ligand-bound receptors in the clathrin domains (*R*_c,S_) and relatively few not clathrin localized (*R*_L,S_). The ligand-bound receptor concentration in clathrin domains *R*_c,S_ remains largely unchanged, with the difference between signaling population and its composition at different clathrin entry rates *k*_+c_ mostly due to changes to the ligand-bound nonclathrin localized receptors *R*_L,S_. Changes to the clathrin entry rate have limited effect on the receptor concentration in clathrin domains because internalization removes receptors in clathrin domains from the cell surface. For the slow rate of background internalization *k*_bi_ and appreciable ligand binding, receptor internalization that occurs from clathrin domains must balance the fixed rate of receptor arrival on the cell surface for the receptor population to reach a steady state.

Figure 3A shows that changing model parameters can shift the magnitude of signaling (level at which signaling saturates) and the ligand binding rate at which signaling rises (ligand binding rate at which the level of signaling receptors reaches half of its maximum). Figure 3C shows how the signaling magnitude changes as both the uniform and localized model parameters are varied from the estimated values. Low internalization and clathrin entry rates and high clathrin exit rates increase signaling, and vice versa, as changes in these parameters affect how many receptors remain on the cell surface. Tetraspanin entry and exit and ligand unbinding do not affect the saturated level of signaling receptors, as changes to these parameters have no effect at sufficiently high ligand binding rates.

Figures 3D–F show how individually varying parameter values affects the ligand binding rate at which concentration of signaling receptors rises for both the uniform and localized models. For the uniform model, lower internalization rate *k*_i_ or clathrin entry rate *k*_+c_ and higher background internalization rate *k*_bi_, clathrin exit rate *k*_-c_, and ligand unbinding rate *k*_-L_ increase the ligand binding rate at which signaling rises (Fig. 3E). Lower internalization rate and clathrin entry reduce internalization, leading to more receptors, particularly ligand-bound and clathrin-localized receptors, on the cell surface. Higher background internalization reduces the receptor concentration available for ligand binding, more clathrin exit reduces internalization, and faster ligand unbinding reduces ligand bound receptors. The effect of changing the shared parameters is similar for the localized ligand binding model (Fig. 3D). Across the parameters shared by both models, Fig. 3F shows that the localized model consistently requires about a threefold higher ligand binding rate (i.e. a threefold higher ligand concentration) for the onset of signaling.

Figure 3D also shows for the localized model that decreasing the tetraspanin entry rate *k*_+T_ or increasing the exit rate *k*_-T_ increases the ligand binding rate for signaling onset. However, unlike the other parameters, *k*_+T_ and *k*_-T_ are not shared with the uniform model. Control of the ligand binding rate for the onset of signaling by the tetraspanin entry and exit rates represents an independent mode of adjusting signaling that is not present in the uniform binding model. Notably, this adjustment of the ligand binding rate for signaling onset via tetraspanin entry and exit rates does not change the maximum level of signaling receptors (Fig. 3C).

The fraction of surface receptors in the signaling states are

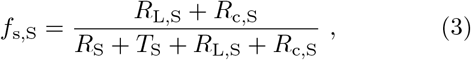

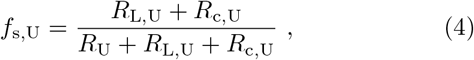

for the spatially localized and uniform ligand binding models, respectively. Figures 3G–I show the changes in the ligand binding rate at which the fraction of surface receptors in the signaling states rises.

The ligand concentration at which the concentration of receptors in signaling states rises (Figs. 3D–F) is much lower than the ligand concentration at which the fraction of receptors in signaling states rises (Figs. 3G–I). The fraction of receptors in signaling states rises at a higher ligand concentration than the concentration of signaling receptors because at higher ligand concentration there is more receptor internalization. At the lower ligand concentrations with substantial concentration of signaling receptors, most receptors do not become ligand bound and localize to clathrin, such that receptor internalization and the decrease in total surface receptor concentration are modest. As the ligand concentration rises further, more receptors become ligand bound and localize to clathrin, causing substantial internalization that decreases the total surface receptor population and leads the receptors in signaling states to be a substantial fraction of the total surface receptor population. The lower total receptor levels on the surface with increased ligand binding rate are shown in the insets of Figs. 3G and 3H for the localized and uniform models, respectively.

For the uniform model, *k*_+L_ ≃ 5 × 10^−2^ s^−1^ for the fraction of receptors in signaling states to rise in Fig. 3H compares to the much lower *k*_+L_ ≃ 6 × 10^−4^ s^−1^ for the concentration of signaling receptors to rise in Fig. 3E. For the localized model, *k*_+L_ ≃ 2 × 10^−1^ s^−1^ for fraction of receptors in signaling states to rise in Fig. 3G compares to the much lower *k*_+L_ ≃ 10^−3^ s^−1^ for signaling receptor concentration to rise in Fig. 3D. For both the uniform and localized models the signaling receptor fraction rises at an approximately hundred-fold higher ligand binding rate (ligand concentration) compared to the signaling receptor concentration. The increase in the signaling receptor concentration as the ligand binding rate rises is curtailed because the surface receptor population becomes severely depleted, such that further increase in the signaling receptor concentration is limited. Only once the surface receptor population is depleted does the signaling receptor fraction substantially rise.

The low physiological EGF concentration of approximately 0.5 ng/mL corresponds to a ligand binding rate of *k*_+L_ = 10^−2^ s^−1^, which is between the relatively low ligand binding rate at which the receptor signaling concentration rises (*k*_+L_ ≃ 10^−3^ s^−1^) and the relatively high ligand binding rate at which the signaling receptor fraction rises (*k*_+L_ ≃ 10^−1^ s^−1^) for the localized model. The low physiological concentration thus corresponds to a ligand binding rate with an elevated level of receptors in signaling states but not an elevated fraction of receptors in signaling states.

The localized model requires an approximately six-fold higher ligand binding rate to reach half of the maximum signaling receptor fraction compared to the uniform model (Fig. 3I). There is less than a factor of two change in ratio of the ligand binding rate for localized to uniform models from up to two orders of magnitude change in parameter values, as measured by the fraction of receptors in signaling states (Fig. 3I).

We also consider by what factor the ligand binding rate must be multiplied to go from low to high signaling, representing the range of ligand binding rates (and ligand concentration) over which signaling can be adjusted (Fig. S2A). We find, denoting low signaling as 10% of the maximum and high signaling as 90% of the maximum, that the range (ratio of *k*_+L_ at high signaling to *k*_+L_ at low signaling) changes little with parameters or between the localized and uniform models (Fig. S2B and S2C).

### Transient signaling

We now focus on transient signaling, with receptor populations changing with time following a sudden change to ligand stimulation. EGF receptor experiments often stimulate the cell with an increase in ligand concentration [37–40] and measure the resulting changes to the receptor population or the signaling response. We similarly began our model receptor populations in steady state for a ligand binding rate of zero before increasing to a non-zero ligand binding rate and allowing the receptor populations to change.

Following a sudden increase in ligand binding rate, the signaling receptor states that are ligand bound outside of and within clathrin domains, the sum of *R*_L_ and *R*_c_, rapidly increase and then gradually decrease, forming an asymmetric peak, with the maximum reached in tens of seconds (Fig. 4A), consistent with experimental measurements [41]. Following the peak, the concentration of receptors in signaling states drops to a new steady state concentration, which is higher than before the increase in the ligand binding rate. We define two quantities to evaluate how the model parameters control these peaks (see Fig. 4A). Peak height is the difference between the concentration at the top of the peak and the concentration at the steady state prior to the ligand binding rate increase. Peak duration is the time spent after the peak maximum in the higher 90% of the concentration range between peak maximum and the new steady state concentration.

**FIG. 4.**
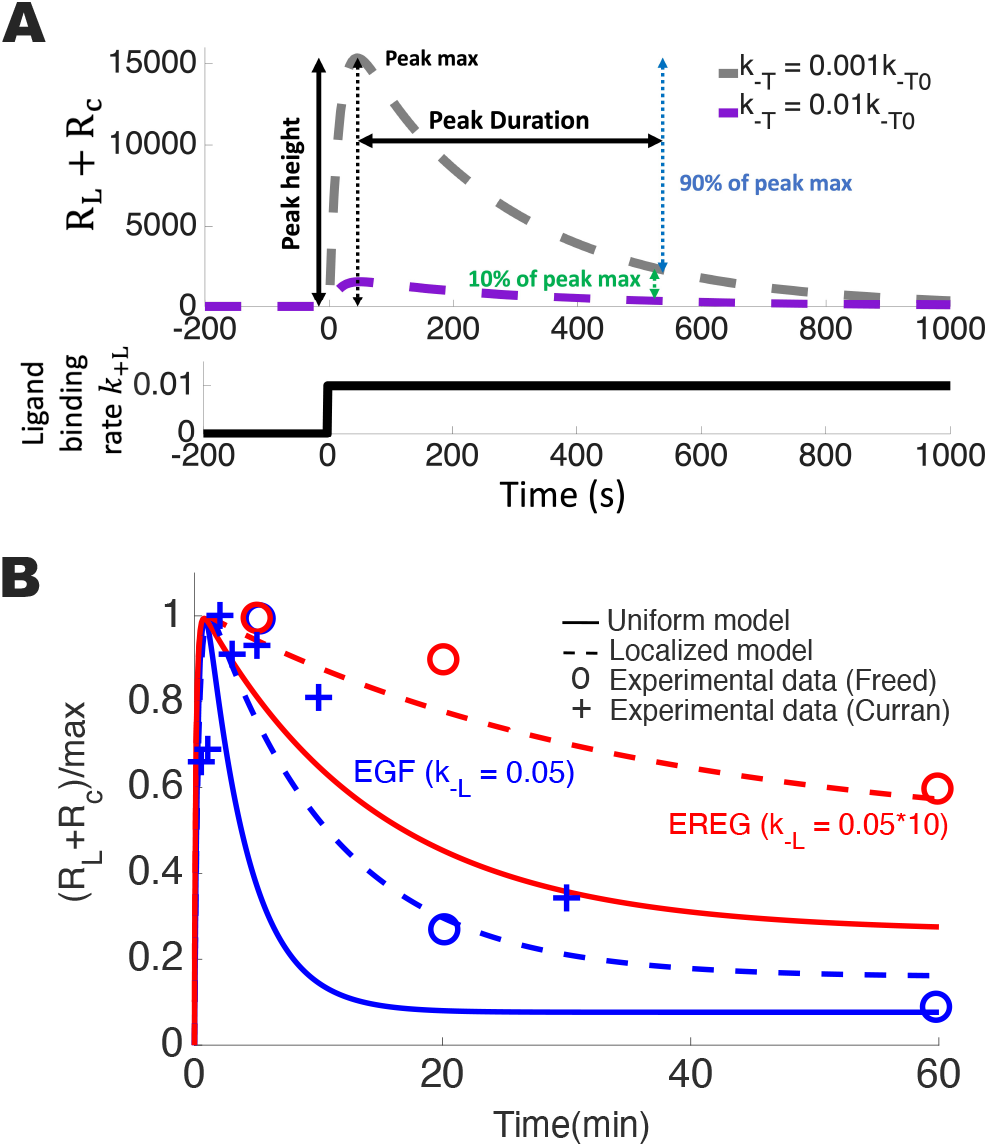
Transient EGFR signaling peaks. (A) Top is the total concentration of EGFR signaling states *R*_L_ + *R*_c_ and bottom is the ligand binding rate *k*_+L_ (units of s^−1^), both vs time. At time of zero the ligand binding rate increases from zero to 0.01 s^−1^. Peak height and duration, which quantify peak behavior in Fig. 5, are indicated, with duration from time of peak maximum until time when concentration has decreased through 90% of the drop to the new steady state. Line color in top part indicates tetraspanin domain exit rate *k*_-T_ with *k*_-T0_ the estimated rate in Table I. (B) Total normalized (divided by maximum) concentration of EGFR signaling states *R*_L_ +*R*_c_ vs time. Curves indicate model data: solid is the uniform model and dashed is the localized model, and blue is a ligand unbinding rate corresponding to EGF ligand (*k*_-L_ = 0.05 s^−1^) and red is corresponding to EREG ligand (*k*_-L_ = 0.5 s^−1^). Points indicate experimental phosphorylation measurement data, with blue for EGF and red for EREG, and circles from Freed et al [37] and crosses from Curran et al [40].

Figure 4A shows that the peak height and duration are affected by model parameters such as the tetraspanin domain exit rate *k*_-T_. Figure 4B shows the transient peaks for two ligand types, corresponding to EGF and EREG, which have binding rates that differ by a factor of ten [12]. Figure 4B also shows experimental signaling data for sudden changes in EGF and EREG concentration [37, 40]. Consistent with experimental data analysis, in Fig. 4B the maxima of the simulated peaks are normalized to one. For the experimental data and both the localized and uniform models, the shorter EREG binding lifetime compared to EGF leads the peak from EREG stimulation to be of much longer duration than the peak from EGF stimulation. This longer-duration peak for shorter binding lifetime is because a receptor with a bound ligand with a shorter binding lifetime is less likely to be internalized [12] and thus more slowly depletes the receptor population on the cell surface. The substantial receptor population that remains for a longer time period allows the signaling to be sustained. In contrast, the longer EGF binding lifetime causes more receptors to be internalized, leaving fewer receptors on the cell surface to continue signaling.

The localized model has peaks of longer duration in comparison to the uniform model (Fig. 4B). This is similar to the shift of steady state signaling onset to higher ligand concentration for the spatially localized model compared to the uniform model in Fig. 3. The peaks from both EGF and EREG stimulation for the spatially localized model also approximately recapitulate experimental data [37, 40] (Fig. 4B), while the uniform model has peaks of substantially shorter duration compared to the experimental data.

In Fig. 5 the peak height and peak duration are systematically explored for the localized and uniform models for an increase from zero ligand binding to a ligand binding rate of 0.01 s^−1^. Figures 5A and 5B show that for both the spatially localized and uniform binding models higher (lower) background internalization rate *k*_bi_ substantially decreases (increases) the peak height, as *k*_bi_ controls the number of receptors remaining on the cell surface. Higher ligand unbinding rate *k*_-L_ substantially decreases and lower *k*_-L_ modestly increases the peak height, as faster ligand unbinding leaves few receptors in the signaling states *R*_L_ and *R*_c_, while for slower ligand unbinding the signaling is less affected if the ligand unbinding becomes slower. Higher (lower) internalization rate *k*_i_ modestly decreases (increases) the peak height, as internalization removes receptors from the cell surface.

**FIG. 5.**
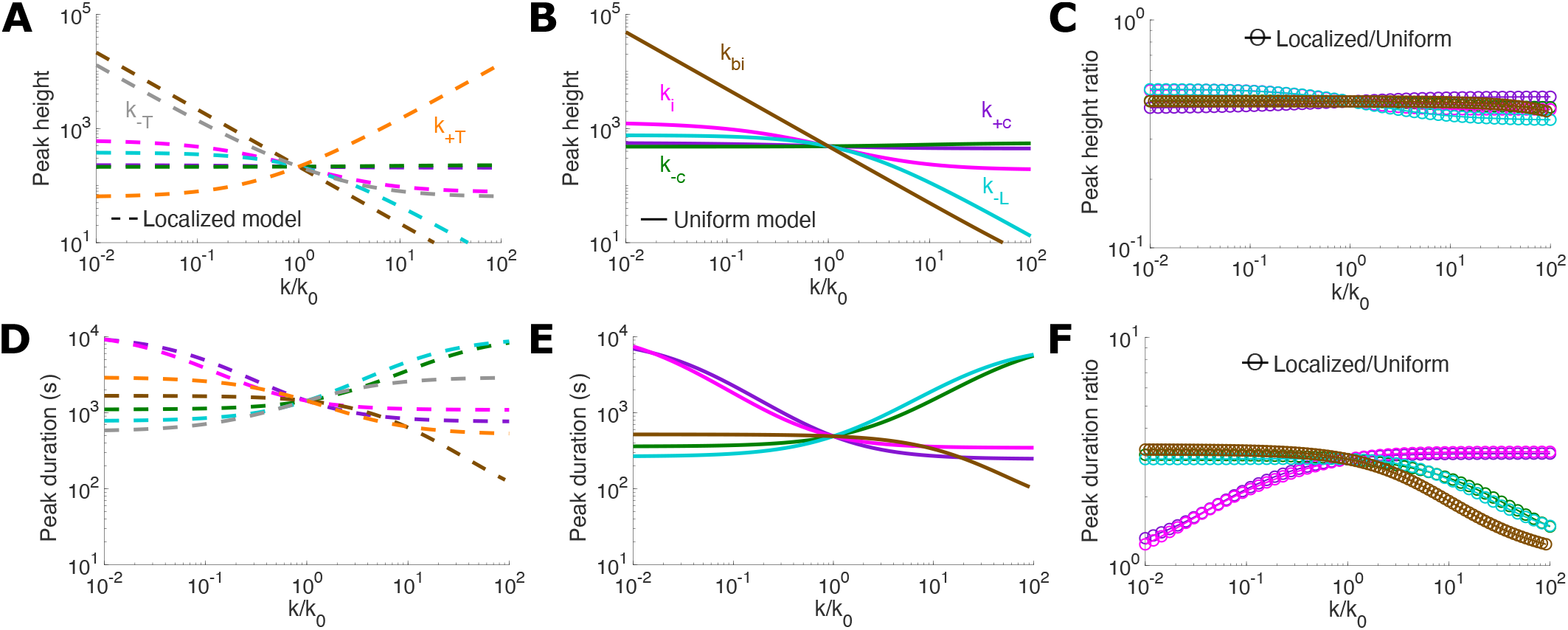
Transient EGFR signaling peak height and duration. (A-B) Peak height as parameters are individually varied for the (A) localized model and (B) uniform model. (C) Ratio, of the localized model to the uniform model, of the peak height, as the shared parameters are individually varied. (D-E) Peak duration (illustrated in Fig. 4A) as parameters are individually varied for the (D) localized model and (E) uniform model. (F) Ratio, of the localized model to the uniform model, of the peak duration, as the shared parameters are individually varied. In all panels, which parameter is varied is indicated by curve color, with estimated rates *k*_0_ from Table I. Curve color labels in panel B apply to all panels, and labels in panel A also apply to panel D. All parameter values are those listed in Table I unless otherwise specified.

Figure 5C shows that the peak height for the localized model is consistently approximately half that of the uniform model as the shared parameter values are varied. Figure 5A shows that beyond the parameters shared by the spatially localized and uniform binding models, changes to the tetraspanin domain entry rate *k*_+T_ and exit rate *k*_-T_ have the largest effects on the peak height, with higher *k*_+T_ increasing and lower *k*_+T_ decreasing the peak height, and vice versa for changes to *k*_-T_. Tetraspanin entry and exit rates represent an independent mode of adjusting the peak height that is not present in the uniform binding model.

Figures 5D and 5E show that adjustments to individual shared parameters (up to two orders of magnitude) for both the localized and uniform models change the peak duration by up to an order of magnitude. Increasing the ligand unbinding rate *k*_-L_ and clathrin exit rate *k*_-c_ and decreasing the internalization rate *k*_i_ and clathrin entry rate *k*_+c_ increase the peak duration because the reduced internalization from clathrin domains more slowly decreases the receptors on the cell surface. Increasing the background internalization rate *k*_bi_ more substantially decreases the peak duration as there is increased removal of receptors from the cell surface.

Figure 5F shows that the peak duration for the localized model is approximately threefold longer than for the uniform model. The peak duration ratio decreases to one for low *k*_i_ or *k*_c_ and high *k*_bi_, *k*_-L_, and *k*_-c_. Thus across much of parameter space the peak duration is longer for the spatially localized binding model compared to the uniform binding model.

As with the peak height in Fig. 5A, the effect of tetraspanin entry and exit rates on peak duration shown in Fig. 5D indicates that the tetraspanin entry and exit rates are an independent mode of adjusting the peak duration that is not present in the uniform binding model. Decreasing the tetraspanin entry rate *k*_+T_ and increasing the tetraspanin exit rate *k*_-T_ increases the duration of the signaling peak (Fig. 5D), although the effect is smaller than for peak height (Fig. 5A).

Figure 5 demonstrates that adjusting ligand unbinding, clathrin entry and exit, and internalization can affect the peak response to ligand increase for both the spatially localized and uniform binding models. However, spatial localization of enhanced ligand binding flattens (reduces the height) and extends (increases the duration) these peaks. Spatial localization of enhanced binding also introduces entry and exit into the primary ligand binding domain (tetraspanin domains for EGFR), which independently of other receptor transitions can be adjusted to make large changes to peak height and smaller but still substantial changes to peak duration.

Figure 5 shows peak behavior for ligand binding rate of zero increased to 0.01 s^−1^, corresponding to a relatively low physiological EGF concentration of 0.5 ng/mL. For an increase from a ligand binding rate of zero to 0.1 s^−1^ (tenfold higher final *k*_+L_ than Fig. 5), the peak behavior is similar, with a somewhat higher and shorter duration peak (see Fig. S1).

Following the increase in the ligand binding rate, the fraction of receptors in signaling states (*R*_L_ + *R*_c_) does not form a substantial peak, but rather increases to a new steady state (Fig. 6A). The fraction of receptors in each state reaches a steady state at the same time (Fig. 6A inset) as the receptors are depleted from the surface. Figure 6B shows the transition duration, the time for the fraction of receptors in signaling states to reach 90% of the new steady state, for the localized model as the parameter values are varied. Across most of the parameter range explored, this transition time is tens of seconds, with the transition time of approximately 40 seconds for the estimated parameter values. Localized and uniform models have quite similar transition times (Fig. 6C), with times for the models remaining within a factor of 1.5 across parameter values.

**FIG. 6.**
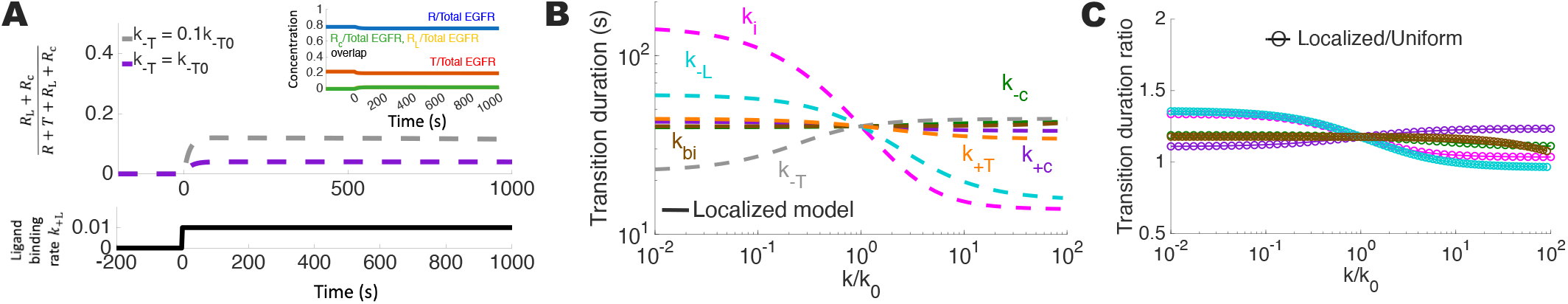
Transient EGFR signaling fraction. (A) Top is the fraction of the total EGFR receptors on the surface that are in the ligand-bound and clathrin localized states (*R*_L_ + *R*_c_) and bottom is the ligand binding rate *k*_+L_, both vs time. At time of zero the ligand binding rate increases from zero to 0.01 s^−1^. Line color in top part indicates tetraspanin domain exit rate *k*_-T_ with *k*_-T0_ the estimated rate in Table I. Inset shows the fraction of surface EGF receptors in each of the four states of the localized model vs time. (B) Transition duration, defined as the time to reach 90% of the new steady state after the ligand binding rate is changed, as individual parameter values are varied for the localized model. Curve color indicates which parameter is varied. (C) Ratio, of the localized model to the uniform model, of the transition duration, as the shared parameters are individually varied. Curve labels for parameters shared by both models in (B) also apply to (C). Estimated rates *k*_0_ are listed in Table I. All parameter values are those listed in Table I unless otherwise specified.

## DISCUSSION

Recent experimental work showed ligand binding to EGF receptors is substantially enhanced for receptors in tetraspanin nanodomains on the cell surface [22]. We have applied quantitative modeling to probe the signaling impact of this spatially localized ligand binding by comparing the behavior of a quantitative model with spatially localized ligand binding to a uniform model with the same ligand binding rate independent of receptor location.

We first recapitulated the experimental finding of unchanging confinement behavior for EGFR as ligand concentration is increased [22] with the spatially localized ligand binding model. The uniform model could not reproduce this experimental result, as the uniform model has confinement rise as the ligand concentration increases. For the localized model the confinement does not change with increasing ligand concentration because receptors move from confinement in tetraspanin domains prior to ligand binding to confinement in clathrin domains following ligand binding. The uniform model, lacking the tetraspanin domain, cannot undergo this change in which domain confines the receptors, and instead the confinement level depends on ligand concentration. Confinement in tetraspanin domains is thus essential for this consistent EGFR confinement across a wide range of ligand concentration. These results suggest that if measurements of other receptors similarly determine that confinement does not change with ligand concentration, then it may indicate multiple confining domains for that receptor. We find that over physiological ligand concentrations ranging from 0.5 ng/mL to 10 ng/mL [22, 30– 33] that EGFR confinement switches from primarily in tetraspanin domains (lower concentrations) to clathrin domains (higher concentrations).

We primarily focused on the levels of EGFR in signaling states, specifically those that are ligand bound outside of or within clathrin domains. For steady state signaling, when ligand concentration is constant in time, our modeling indicates that spatially localized ligand binding increases (compared to uniform binding) the ligand concentration required to induce a certain level of steady state signaling. For the estimated parameters, the ligand concentration required for signaling onset increases by approximately a factor of three, from about 0.02 ng/mL (corresponding to a ligand binding rate of 4 × 10^−4^ s^−1^) for uniform binding to about 0.05 ng/mL (ligand binding rate of 10^−3^ s^−1^) for spatially localized binding. These ligand concentrations are relatively low compared to physiological ligand concentrations of about 0.5 ng/mL [30–32]. The correspondence of the responsive signaling range to these low EGF concentrations suggests that the shift of the responsive EGF concentration range to higher concentrations for the spatially localized binding model (compared to the uniform model) may enable EGF receptors to provide varying levels of signaling output between low and high physiological EGF concentrations. We did not find that the range of ligand concentration over which signaling changes from low to high (ratio of ligand concentrations for high and low signaling) is affected by localized vs uniform ligand binding or parameter values.

For steady-state signaling, our model indicates two groups of parameters. Clathrin domain recruitment and internalization primarily control the magnitude of the signaling response, whereas tetraspanin domain recruitment and ligand unbinding rate control the ligand concentration required to reach a given level of signaling receptors.

We find that there are two distinct signaling regimes with meaningful concentration of signaling receptors. At a sufficiently low ligand concentration, there is a vanishing concentration of signaling receptors. For medium ligand concentration, the concentration of receptors in signaling states reaches a substantial fraction (tens of percent) of the maximum concentration of receptors in signaling states (found at very high ligand concentration), but the receptors in signaling states are a tiny fraction of all surface receptors. For high ligand concentration, although the concentration of signaling receptors has only risen modestly compared to the medium ligand concentration, the receptors in signaling states are a large fraction (tens of percent) of the surface receptor population, which has been decreased by more than an order of magnitude compared to low ligand concentration. The medium ligand concentration and high ligand concentration regimes, defined as the ligand concentrations at which the signaling receptor concentration and signaling receptor fraction reach half their maximum value, respectively, are at distinct ligand concentrations separated by approximately two orders of magnitude. We find that for the spatially localized ligand binding EGFR model the lower end of physiological EGF concentrations (≃ 10^−2^ s^−1^ ligand binding rate, equivalent to 0.5 ng/mL [30–32]) is between the medium (ligand binding rate of 10^−3^ s^−1^, equivalent to 0.05 ng/mL) and high (ligand binding rate of 10^−1^ s^−1^, equivalent to 5 ng/mL) ligand concentration regimes. This suggests that not just the concentration of signaling (ligand-bound and clathrin-confined) EGFR is important for signaling response, but that the fraction of receptors on the surface in a signaling state may be important as well. This aligns with work demonstrating that receptor abundance and ligand-bound fraction are key factors for EGFR signaling [42] and with the relevance attributed to fraction of occupied receptors across receptor types [43, 44].

For transient signaling following a sudden change to ligand concentration, we find the signaling receptor states form an asymmetric peak that rises quickly and then falls more slowly, trending to a new steady state. We examined the magnitude and duration of the transient peak in the concentration of signaling receptors. With the localized ligand binding model the peak is extended to about three-fold the duration as with the uniform model. With the localized model, the duration, which represents the time to approach the new steady state, is approximately 10^3^ seconds (about 15 to 20 minutes) for the estimated parameters with EGF as the ligand and 5 × 10^3^ seconds (about an hour) for with EREG (10 × faster unbinding compared to EGF) as the ligand, both with the localized model. These durations are consistent with experimental data measuring signaling following a change in EGF receptor ligand (EGF and EREG) stimulation, for which EGF stimulation induces a peak in signaling receptors that decays in about 20 minutes, while EREG stimulation induces a peak that is sustained, retaining most of the peak height after an hour [37, 40]. This sustained signaling from EREG stimulation, in contrast with the transient peak that quickly returns to a similar steady state induced by EGF, may be one aspect of signaling that allows a cell to distinguish between signals. The persistence of signaling as an identifier of signal type aligns with the broad importance of temporal patterns for cell signaling [45, 46], including for EGFR [47, 48].

The duration of the peak of receptors in signaling states following an increase in ligand concentration informs the timing of measurements in experiments. Measurements taken at times before the steady state is approached (i.e., before about 10 minutes following the change in ligand concentration for EGF) are sampling from the transient response, with elevated signaling receptor levels, rather than a new steady state. Spatially localized ligand binding reduces the magnitude (by about a factor of two) and extends the duration (by about a factor of three) of the transient increase in EGF receptors in signaling states, compared to uniform binding.

Because increasing the ligand concentration causes depletion of the receptor level on the cell surface, the fraction of receptors in signaling states does not rise and then meaningfully fall (i.e., form a substantial peak) following increase in the ligand concentration. Instead the fraction of surface receptors in a signaling state goes almost directly to the new steady state in a much shorter time, requiring about a minute for the fraction of receptors in signaling states vs. 15 – 20 minutes for the concentration of receptors in signaling states to reach the new steady state. This difference is because the fraction of receptors in signaling states is nearly at the steady state when the peak of the concentration of receptors in signaling states is reached. Following the peak, the fraction of receptors in each state changes little, instead the concentrations in each EGFR state decrease together as receptors are depleted from the cell surface. The signaling receptor concentration slowly decreases from the peak to the new steady state.

For our model, receptors are severely depleted from the cell surface for higher ligand concentrations. Receptor level depletion is important for the fraction of receptors in signaling states, as much of the increase in this fraction is due to the decrease in total receptor levels in addition to the increase in the level of signaling receptors. The depletion our model predicts is more than seen experimentally (about 50% depletion followed by partial recovery [49]), pointing to future work to incorporate EGFR recycling [49] into the model. Our model does not include EGFR oligomerization, which plays a role in EGFR signaling [14, 17, 50, 51], consideration of which also indicates a fruitful future direction.

We find that compared to uniform ligand binding, spatially localized ligand binding in tetraspanin domains increases the ligand concentration for signaling onset for steady state signaling, and decreases the peak height and increases the peak duration for the transient response to ligand concentration increase. Adjustment of the processes shared by both the spatially localized and uniform ligand binding models (ligand unbinding, internalization, clathrin domain entry and exit) yields very similar changes to the level of signaling receptors, for both both steady state and transient scenarios. This indicates that introduction of spatially localized ligand binding does not change how these processes affect signaling. In contrast, signaling changes caused by adjusting entry and exit into tetraspanin domains for the spatially localized ligand binding model represent an independent regulatory mode for both steady state and transient EGF receptor signaling compared to what is enabled by the uniform ligand binding model.

By requiring a stronger signal for a given steady state response, decreasing entry into and increasing exit from tetraspanin domains may limit the response of the cell to EGF receptor signals overall, and increasing entry and decreasing exit may increase the responsiveness of the cell to all EGF receptor signals. Similarly, decreasing entry and increasing exit could extend the response to changing signals, and increasing entry and decreasing exit could shorten the response period, adjust downstream signaling aspects regulated by signal duration.

Rates of EGFR receptor entry and exit into tetraspanin nanodomains may be altered by modifications to the protein states and protein populations in tetraspanin domains, similar to protein recruitment to clathrin domains [52]. Furthermore, the size and number of tetraspanin nanodomains (which may be controlled by tetraspanin protein levels) are expected to affect the speed of diffusive search for these domains [53]. EGFR recruitment to tetraspanin nanodomains could also be impacted indirectly, for example via cholesterol levels which modify membrane fluidity [54, 55] and affect the diffusivity of embedded proteins [56] and diffusive search times — accordingly factors such as cholesterol levels, which can be altered in cancer cells [57, 58], could alter tetraspanin entry and exit rates.

## CONCLUSION

By applying quantitative modeling we have shown that enhanced ligand binding in tetraspanin domains impacts the level and required stimulation for EGF receptor signaling, as well as the duration of transient signaling in response to stimulation changes. Similar changes may occur for other receptors with enhanced activation when localized to specific domains on the cell surface.

## APPENDIX

### Model differential equations and steady-state solutions

The differential equations that describe the spatially localized ligand binding model (Fig. 1A) are

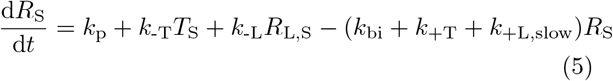

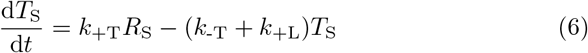

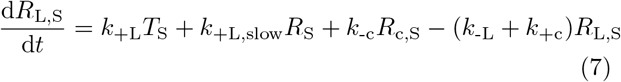

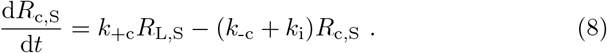

The steady state solution, determined by setting all derivatives in Eqs. 5 to 8 to zero, is

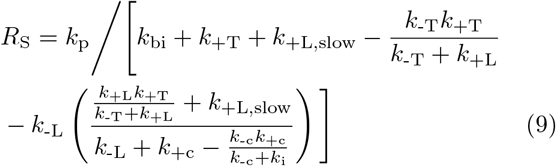

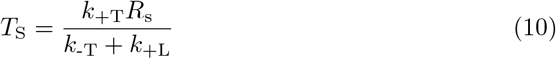

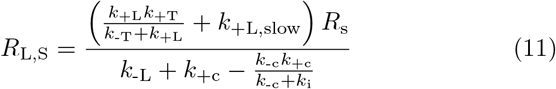

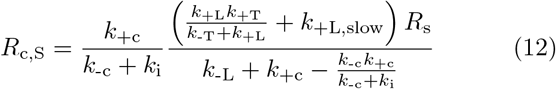

The differential equations that describe the uniform ligand binding model (Fig. 1B) are

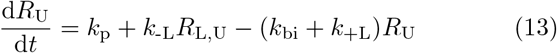

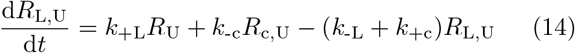

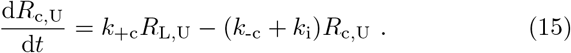

The steady state solution, determined by setting all derivatives in Eqs. 13 to 15 to zero, is

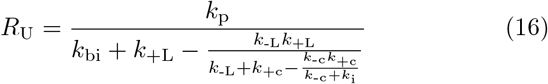

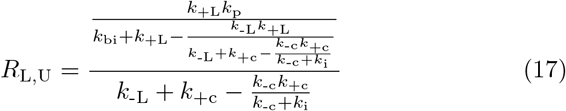

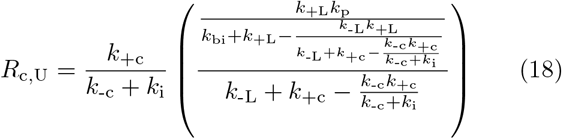

### Kinetic model parameter estimation

We use an estimated background internalization rate of *k*_bi_ = 3 × 10^−4^ s, as constitutive EGF receptor internalization has been measured as 0.02 – 0.05 min^−1^ (3 × 10^−4^ −8 × 10^−4^ s^−1^) [59, 60]. The ligand unbinding rate *k*_-L_ is estimated as 0.05 s^−1^ for EGF and 0.5 s^−1^ for EREG [12]. The clathrin domain entry rate is estimated as *k*_+c_ = 0.1 s^−1^, the clathrin domain exit rate as *k*_-c_ = 0.05 s^−1^, and the internalization of clathrin-localized receptors as *k*_i_ = 0.05 s^−1^ [12].

We estimate the ligand binding rate by assuming that the process of ligand binding is diffusion-limited, that a ligand that reaches a receptor will bind to the receptor. For a fixed concentration *ρ* of ligands of diffusivity *D* far from a spherical cell of radius *R* with *N* absorbing receptors of radius *s* on the surface, the diffusion-limited rate *k*_+L_ of ligand to find each receptor is [53]

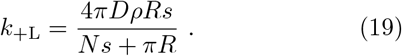

With typical physiological EGF ligand concentration measured as 200 pg/mL [30], 500 pg/mL [31], and 700 pg/mL [32] and higher physiological concentrations of 5 – 10 ng/mL noted [22], a 0.5 ng/mL concentration corresponds to approximately (EGF is 600 kDa [61]) 0.5 EGF molecules/*µ*m^3^. 10^4^ to 10^5^ EGF receptors are present on the cell surface for some cell types [62]. EGF diffusivity has been measured as 166 *µ*m^2^*/*s in dilute agarose gel and in the rat brain extracellular volume as 51.8 *µ*m^2^*/*s [63], and accordingly we estimate the EGF diffusivity as 5 *µ*m^2^*/*s to 50 *µ*m^2^*/*s. With EGF receptors approximately 2.5 nm in radius [62], we estimate that the ligand binding site is approximately 1 nm in radius, and that the cell is 20 *µ*m in radius. Then *k*_+L_ ranges from 0.004 s^−1^ to 0.08 s^−1^, of which a intermediate value is approximately 0.01 s^−1^ corresponding to a ligand concentration of 0.5 ng/mL.

Experimental observations show most of the decay in surface EGFR levels following exposure to 10 nM EGF, down to about 40% of pre-exposure receptors levels, occurs over a time period of about five minutes [49]. We can represent this decay as following *e*^−*t/τ*^ = 0.4, with *t* = 300 s and *τ* the time period for the underlying internalization process to occur once, so *τ* = −*t/* log(0.4) = (300 s)*/* log(0.4) ≃ 330 s. The underlying process involves tetraspanin domain entry (period *T*_tetra_), ligand binding (*T*_bind_), clathrin entry (*T*_clathenter_), clathrin exit if not internalized (*T*_clathexit_), and internalization from clathrin domains (*T*_bind_). As EGF has about a 29% probability of resulting in internalization for each binding event [12] and only about 50% of receptors will be internalized from clathrin domains and 50% will exit and re-enter, then *τ* = (*T*_tetra_ + *T*_bind_ + 2*T*_clathenter_ + *T*_clathexit_ + *T*_int_)*/*0.29. With 10 nM EGF equivalent to approximately 60 ng/mL and a ligand binding rate of 1 s^−1^ then *T*_bind_ = 1 s, *T*_clathenter_ = 1*/k*_+c_ = 10 s, *T*_clathexit_ = 1*/k*_-c_ = 20 s, and *T*_int_ = 1*/k*_i_ = 20 s. Thus *T*_tetra_ = 0.29*τ* − *T*_bind_ − 2*T*_clathenter_ − *T*_clathexit_ − *T*_int_ = 0.29(260 s) − 1 s − 20 s − 20 s − 20 s ≃ 30 s. *k*_+T_ = 1*/T*_tetra_ ≃ 0.033 s ^1^. In conditions with no stimulating EGF or other ligand, approximately 20% of EGFR are in tetraspanin domains [22], i.e. *T/*(*R* + *T* ) = 0.2 or *R/T* = 4. In this scenario entry and exit from clathrin domains is balanced, *k*_+T_*R* = *k*_-T_*T*, so *k*_-T_ = *k*_+T_*R/T*≃ 0.12 s^−1^.

### Additional results

Figure 7 shows how the peak height and duration change in response to an increase in the ligand binding rate from zero to 0.1 s^−1^, similar to Fig. 5, which is for an increase to a ligand binding rate of 0.01 s^−1^. Figure 8 shows how the ratio of ligand binding rate that provides a high signaling level (90% of the maximum) to the ligand binding rate that provides a low signaling level (10%) of maximum changes between the localized and uniform models, and with parameter changes.

**FIG. 7.**
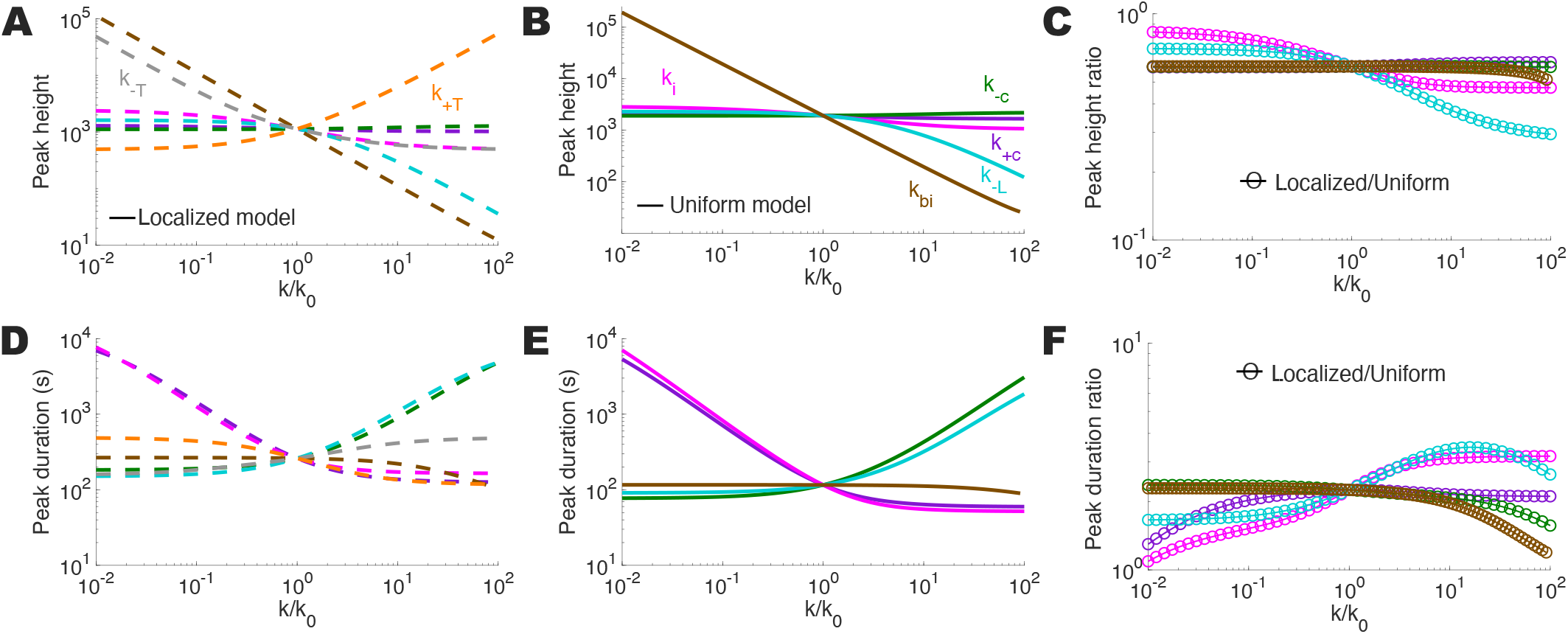
Transient EGFR signaling peak height and duration for higher ligand binding rate after increase (0.1 s^−1^ compared to 0.01 s^−1^ in Fig. 5). (A-B) Peak height as parameters are individually varied for the (A) localized model and (B) uniform model. (C) Ratio, of the localized model to the uniform model, of the peak height, as the shared parameters are individually varied. (D-E) Peak duration (illustrated in Fig. 4A) as parameters are individually varied for the (D) localized model and (E) uniform model. (F) Ratio, of the localized model to the uniform model, of the peak duration, as the shared parameters are individually varied. In all panels, which parameter is varied is indicated by curve color, with estimated rates *k*_0_ from Table 1. Curve color labels in panel B apply to all panels, and labels in panel A also apply to panel D. All parameter values are those listed in Table 1 unless otherwise specified.

**FIG. 8.**
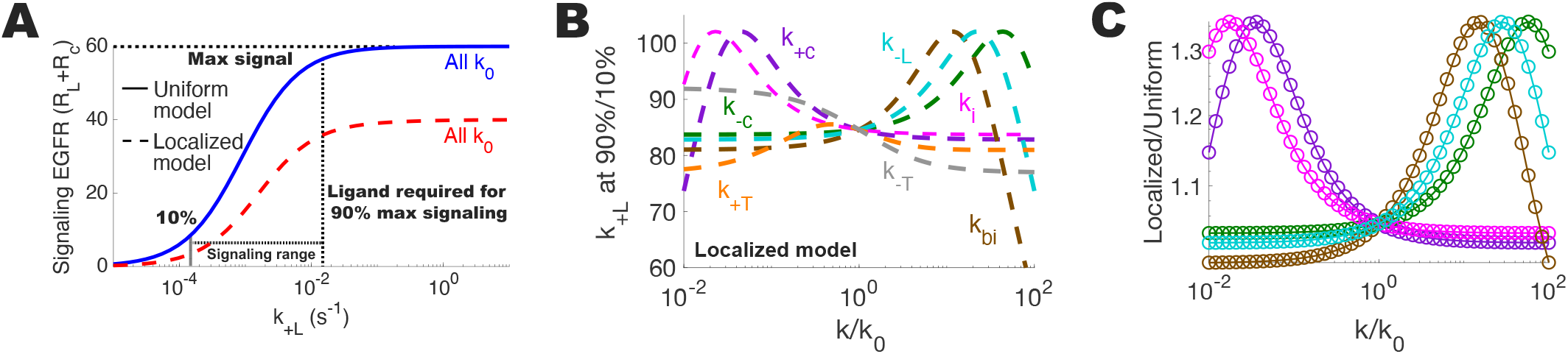
Steady state signaling range at 10% and 90%. (A) Signaling EGF receptor (*R*_L_ + *R*_c_) concentration as the ligand binding rate is varied. Parameter values indicated by curve color, with ‘All *k*_0_’ indicating that all model parameters are the estimated parameter values from Table 1. The horizontal dotted black line indicates the maximum level of signaling receptors and the two vertical dotted black line indicates the ligand binding rate *k*_+L_ at which the level of signaling receptors is 10% and 90% the maximum level of signaling receptors, both for the uniform model (blue solid curve). (B) The ratio of ligand required for high (90% of maximum *R*_L_ + *R*_c_) and low (10% of maximum *R*_L_ + *R*_c_) signaling for the localized model as model parameters are varied. *k/k*_0_ indicates the ratio of the parameter value and the estimated parameter value listed in Table 1. C) The ratio of high to low signaling (as shown in B) for the localized model over the uniform model. The uniform model is nearly constant, so the ratio just scales the values from (B), leaving the curve shape unchanged.

